# A family of contact-dependent nuclease effectors contain an exchangeable, species-identifying domain

**DOI:** 10.1101/2020.02.20.956912

**Authors:** Denise Sirias, Daniel R. Utter, Karine A. Gibbs

## Abstract

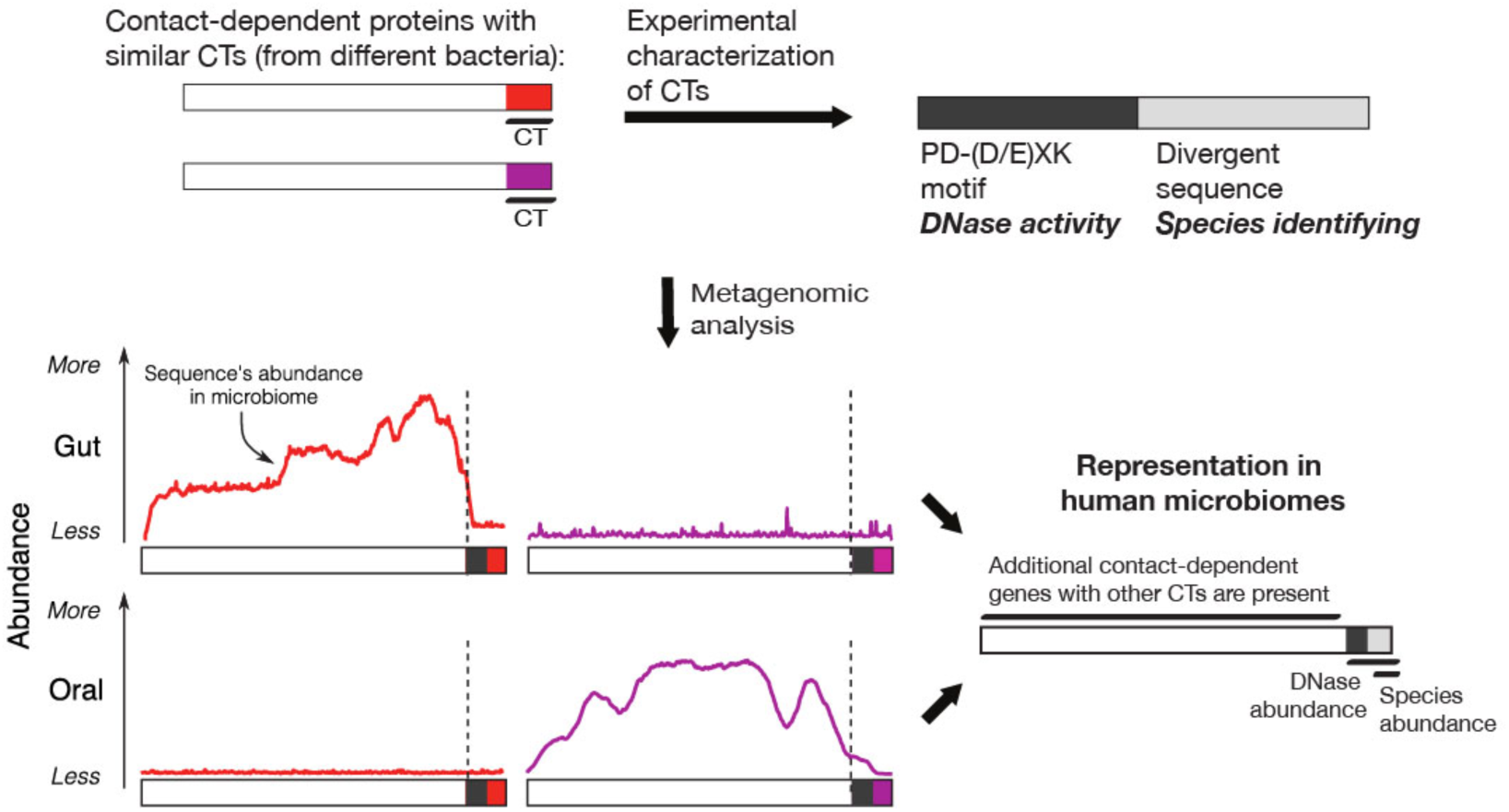

In mixed-species communities, bacteria can deploy contact-dependent effectors to compete with other organisms, often directly injecting these proteins into neighboring cells. One current hypothesis is that the entire protein contains information specific for a single species; emergence of new effectors comes from acquiring genes. Here we have characterized a family of DNA-degrading effectors that are nucleases which cause death. Like other families of chimeric nucleases, these effectors contain two domains. One is a PD-(D/E)XK-containing domain necessary for DNA cleavage. The other domain, which does not contain known DNA-binding structures, encodes species-identifying information. We capitalized on the species-identifying domain to differentiate among low-abundance species, as well as to reveal domain architectures within these proteins, in human gut and oral microbiomes. Emerging are questions about how low-abundance strains use effectors for survival and how strain-identifying effectors evolve.

## Introduction

Bacteria within communities compete for resources. One strategy is to inject lethal effectors into adjacent cells; clonal siblings survive by neutralizing the effector while all other cells die (*1–3*). Such bacterial effectors, which can act in many ways such as disrupt cell membranes and degrade proteins or nucleic acid (*4*), are particularly useful for bacteria in gut microbiomes and other mixed communities. Individual strains of a single species can encode different effectors, each with different structures (*5–7*). Abundant species use their specific effectors to dominate in a mixed-species community (*8, 9*). Yet low-abundance bacteria can remain successful residents within mixed-species communities that have a dominant strain; this mechanism for survival remains unclear. Moreover, strain-specific proteins could provide a genome-encoded mechanism to differentiate among closely related bacteria.

The swarming bacterium *Proteus mirabilis*, a human opportunistic pathogen, is one of the low-abundance bacteria in some human guts. In *P. mirabilis*, the gene *idrD*, encoding a contact-dependent lethal effector, was identified in a screen for self versus non-self recognition (*10*). Strains lacking a functional *idrD* gene are unable to merge colonies with their wild-type parent or dominate in mixed migrating populations (*10*) and possibly in the human host (*11*). IdrD is exported by a Type VI secretion system, allowing for transport through direct physical contact with adjacent cells (*10, 12*). The C-terminal end of IdrD encodes a potential cell-contact dependent effector protein, IdrD-CT.

Polymorphic effectors like IdrD contain regions with known functions: the N-terminal domain aids in the export via Types IV, V, or VI secretion systems (*1, 13, 14*) and the C-terminal domain often contains the lethal effector (*2, 3, 7, 15*). For example, the well-established polymorphic effectors RhsA and RhsB have their enzymatic nuclease domains at the C-terminus; putative proteolytic cleavage results in the smaller effectors, RhsA-CT and RhsB-CT, respectively (*2, 16*). Interestingly, effectors with similar functions, such as degrading DNA, can be of different protein families, each with distinct secondary and tertiary structures. This domain architecture could promote the existence of species-identifying effectors, leading to a hypothesis that the effector, encoded within the entire C-terminal domain, encodes specificity for a species (*17*). However, we found that proteins with similarity to IdrD-CT contained more sequence variation than could be explained by this hypothesis.

The uncharacterized IdrD-CT protein was particularly interesting because secondary and tertiary structure analysis predicted the presence of a PD-(D/E)XK motif, which is found in a broad superfamily of nucleases. Contact-dependent effectors in *Burkholderia pseudomallei* that degrade tRNA (*18*) are an example of such nucleotide-targeting enzymes. Database searches revealed additional proteins with similarity to IdrD-CT, yet none of these proteins belonged to known families of the PD-(D/E)XK nuclease superfamily. Even though proteins containing the PD-(D/E)XK motif are found in animals, plants, and bacteria (*19*), predicting the nucleotide target from the amino acid sequences remains difficult. Finding and characterizing additional families is critical for developing algorithms that could predict function from sequence.

As such, we sought to characterize the function and nucleotide target of the IdrD-CT protein. Through biochemical assays, we found that IdrD-CT, and proteins with similar sequences, form a distinct family of PD-(D/E)XK nuclease proteins. We next investigated whether one could differentiate between low and high abundance bacteria in human datasets by adapting metagenomics methods to find IdrD-CT. Metagenomes are short-read DNA datasets generated from all bacteria within a sampled community or microbiome. An adapted metagenomic analysis revealed that members of the IdrD-CT protein family are rare in human oral and gut microbiomes, even though the full-length encoding gene, *idrD*, is not. Analysis of metagenomes also showed that IdrD-CT proteins have an architecture with two separate domains, one encoding deoxyribonuclease (DNase) activity and the other with species-identifying sequences. IdrD-CT has a conserved cleavage action in the DNase domain and is flexible to amino acid changes, including entire domain replacements, in the species-identifying domain. Species-identifying effectors could provide a means to differentiate among bacterial species within gut and oral microbiome datasets. The domain architecture of the IdrD-CT protein and related proteins raise questions about how strain-specific effectors evolve and how bacteria might deploy these effectors for survival at low abundance.

## Results

We found that the IdrD-CT proteins degrade DNA and form a family of PD-(D/E)XK proteins. *P. mirabilis* strain BB2000 needs the IdrD-CT protein to outcompete in mixed-strain populations (*10*), yet the protein had an unknown activity. We found a potential PD-(D/E)XK domain (1 – 86 amino acids) residing in the N-terminal region. PD-(D/E)XK-containing proteins can function in DNA repair, restriction, replication, or tRNA–intron splicing (*19, 20*). We thus hypothesized that IdrD-CT might be a nucleotide-targeting protein.

If this hypothesis was correct, then IdrD-CT should prevent cell growth. Further, mutation of the PD-(D/E)XK catalytic site should disrupt the protein activity. In support of these predictions, we found that *P. mirabilis* cells engineered to overproduce IdrD-CT had 1000 fewer cells per mL than cells not overproducing this protein (Fig. 1A). An equivalent growth pattern occurs in *Escherichia coli* cells engineered to overproduce the *P. mirabilis* IdrD-CT (Fig. S1A), indicating that the target is common among these bacteria. Three residues in the catalytic site (D, D/E, and K) are essential for nuclease function in PD-(D/E)XK-containing proteins (*21, 22*). We replaced each residue in IdrD-CT (D39, E53, and K55) with an alanine residue individually and all at once. *P. mirabilis* producing these mutant proteins grew to a density equal to that of the negative control lacking IdrD-CT (Fig. 1A), indicating that these residues are essential for preventing cell growth. We concluded that IdrD-CT is a PD-(D/E)XK-containing protein.

**Fig. 1.**
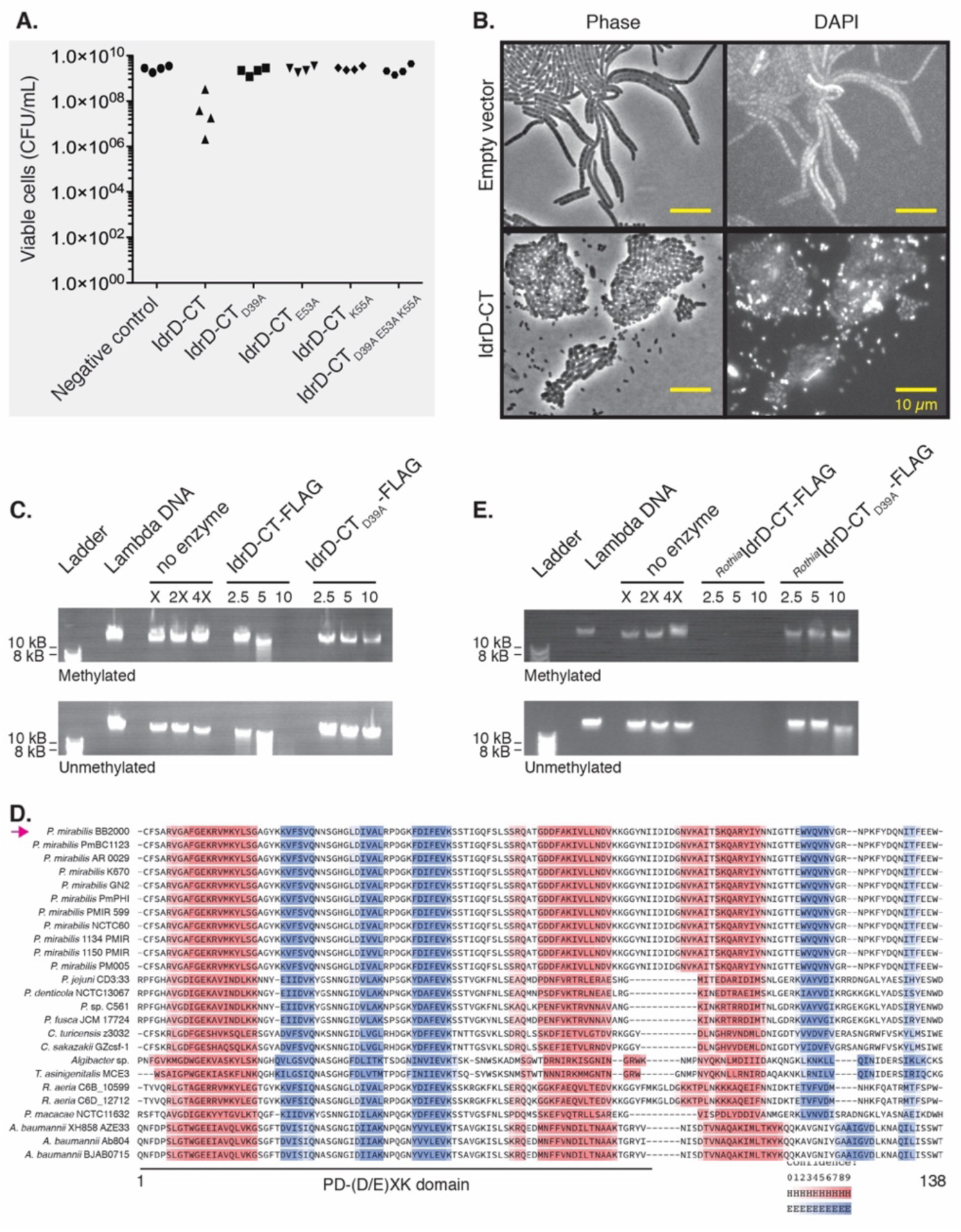
IdrD-CT is a DNA-degrading enzyme in the PD-(D/E)XK superfamily. (A) Displayed are colony-forming units per mL (CFU/mL) for *P. mirabilis* cells after excess production of GFP (negative control), IdrD-CT or mutant variants, as denoted. Protein is produced from a plasmid. (B) The micrographs show *P. mirabilis* grown on a swarm-permissive surface. IdrD-CT is *P. mirabilis* overproducing the wild-type protein from a plasmid. The negative control is *P. mirabilis* with an equivalent empty vector; healthy *P. mirabilis* cells elongate in swarms. Left, Phase. Right, fluorescence of DAPI-stained DNA. (C) IdrD-CT-FLAG or IdrD-CT_D39A_-FLAG (2.5, 5 and 10 ng) were added to methylated and unmethylated lambda DNA (48,502 base pairs, bp), and assayed on an agarose gel with a 2-log DNA ladder. Only samples with IdrD-CT show reduction in DNA. (D) Proteins like IdrD-CT are aligned (using MUSCLE) with predicted secondary structures as colors on top, generated using (*23*). Red are alpha-helices, and blue are beta-sheets; color intensity reflects confidence of the predictions. The PD-(D/E)XK domain is noted. An arrow marks the IdrD-CT from *P. mirabilis* strain BB2000. (E) *^Rothia^*IdrD-CT-FLAG and *^Rothia^*IdrD-CT_D39A_-FLAG (2.5, 5 and 10 ng) was added to lambda DNA and assayed as in (C). Samples with *^Rothia^*IdrD-CT show a reduction in DNA.

Experimental determination of IdrD-CT’s nucleotide target was necessary as predicting the nucleotide target from the amino acid sequence is still difficult for the PD-(D/E)XK nuclease superfamily. Imaging of wild-type *P. mirabilis* producing excess IdrD-CT from a vector showed an accumulation of misshapen cells with frequent absence of DNA (Fig. 1B). By contrast, control samples with no produced protein had characteristic elongated *P. mirabilis* cells; fluorescence associated with DNA was regularly spaced along the length of the cells (Fig. 1B). *E. coli* cells producing IdrD-CT also had a morphology consistent with DNA damage or stress; cells were elongated and had irregular subcellular localizations of DAPI-stained DNA (Fig. S1B). By contrast, *E. coli* cells without IdrD-CT were short with visible DAPI-stained DNA (Fig. S1B). Fluorescence due to DAPI was spaced regularly within each elongated cell of *E. coli* producing the null mutant IdrD-CT_D39A_ (Fig. S1B), indicating that disruption of the predicted active site in IdrD-CT prevents DNA disruption in these cells.

The microscopy results implicated DNA as a likely target for IdrD-CT. To assess this possibility, we made IdrD-CT protein (∼ 17 kDa) and the null mutant protein, IdrD-CT_D39A_, each containing a FLAG epitope tag for detection, in a cell-free extract and then introduced phage lambda DNA. Degradation of lambda DNA occurred in the presence of IdrD-CT, regardless of the DNA methylation state (Fig. 1C, Fig. S1C). However, the null mutant IdrD-CT_D39A_, as well as the negative control in which no IdrD-CT is present, showed little reduction in lambda DNA (Fig. 1C). Across all reactions, no apparent reduction in RNA occurred (Fig. S1C). The IdrD-CT protein, but not the mutant protein, also degraded circular DNA (Fig. S1D). We concluded that the *P. mirabilis* IdrD-CT is capable of degrading both methylated and unmethylated DNA.

To examine whether IdrD-CT was produced in bacteria beyond *P. mirabilis*, we conducted a phylogenetics screen for similar proteins. We identified 23 additional proteins through searches of UniProtKB and the NCBI draft and completed genomes using phmmer (*24, 25*) and tblastn (*26, 27*). Each identified protein has the three conserved catalytic residues (D, D/E, K). Predicted secondary structures are consistent within these predicted proteins (Fig. 1D). None of these IdrD-CT proteins belonged to known PD-(D/E)XK-containing families.

Given the sequence similarity, we queried whether these identified proteins also have DNA-degrading activity. We focused on a *Rothia aeria* protein as *R. aeria* are evolutionarily distant from *P. mirabilis*. *R. aeria* are Gram-positive inhabitants of the normal flora within the human oral cavity and pharynx (*28–30*). Using the cell-free system, we produced the *Rothia* IdrD-CT protein, encoded by gene C6B_10599, and a predicted null mutant *Rothia* IdrD-CT_D39A_. Each protein had a C-terminal FLAG epitope tag for detection. Samples with the *Rothia* IdrD-CT protein showed a loss of lambda DNA regardless of methylation state, while samples containing the *Rothia* IdrD-CT_D39A_ protein or a negative control did not (Fig. 1E, Fig. S1E). There was no observable loss in RNA (Fig. S1E). Therefore, the *Rothia* IdrD-CT protein also degrades DNA.

Additional phylogenetic reconstruction of the identified proteins found representation across the bacterial tree (Fig. S2A). Protein sequences within a specific genus were more related than to those of different genera (Fig. S2A). However, a species tree based on the full-length 16S rRNA gene (Fig. S2B) revealed a discordant topology to the IdrD-CT tree (Fig. S2A). Proteins from evolutionarily distant bacteria appeared to share more similarity in sequence than with more closely related genera. Even with this topology, the *Proteus* and *Rothia* IdrD-CT proteins retained equivalent activities. We concluded that these IdrD-CT proteins, found across bacterial clades, form a distinct family of DNase proteins.

### IdrD-CT is modular and contains an interchangeable species-identifying domain

In the sequence alignments of the identified proteins (Fig. 1D), the C-terminal region contained greater variability in amino acid composition across family members than the N-terminal region. We hypothesized that IdrD-CT proteins have two domains: one containing the PD-(D/E)XK motif and one containing species-identifying information. Therefore, we made independent deletions of each domain in the *P. mirabilis* IdrD-CT protein and assayed for activity. Both deletion proteins resulted in no loss of lambda DNA (Fig. 2A, Fig. S3A). Equivalent truncations of the *Rothia* IdrD-CT protein also showed no DNA degradation (Fig. S3B). Therefore, both the enzymatic and species-identifying domains are essential for IdrD-CT activity.

**Fig. 2.**
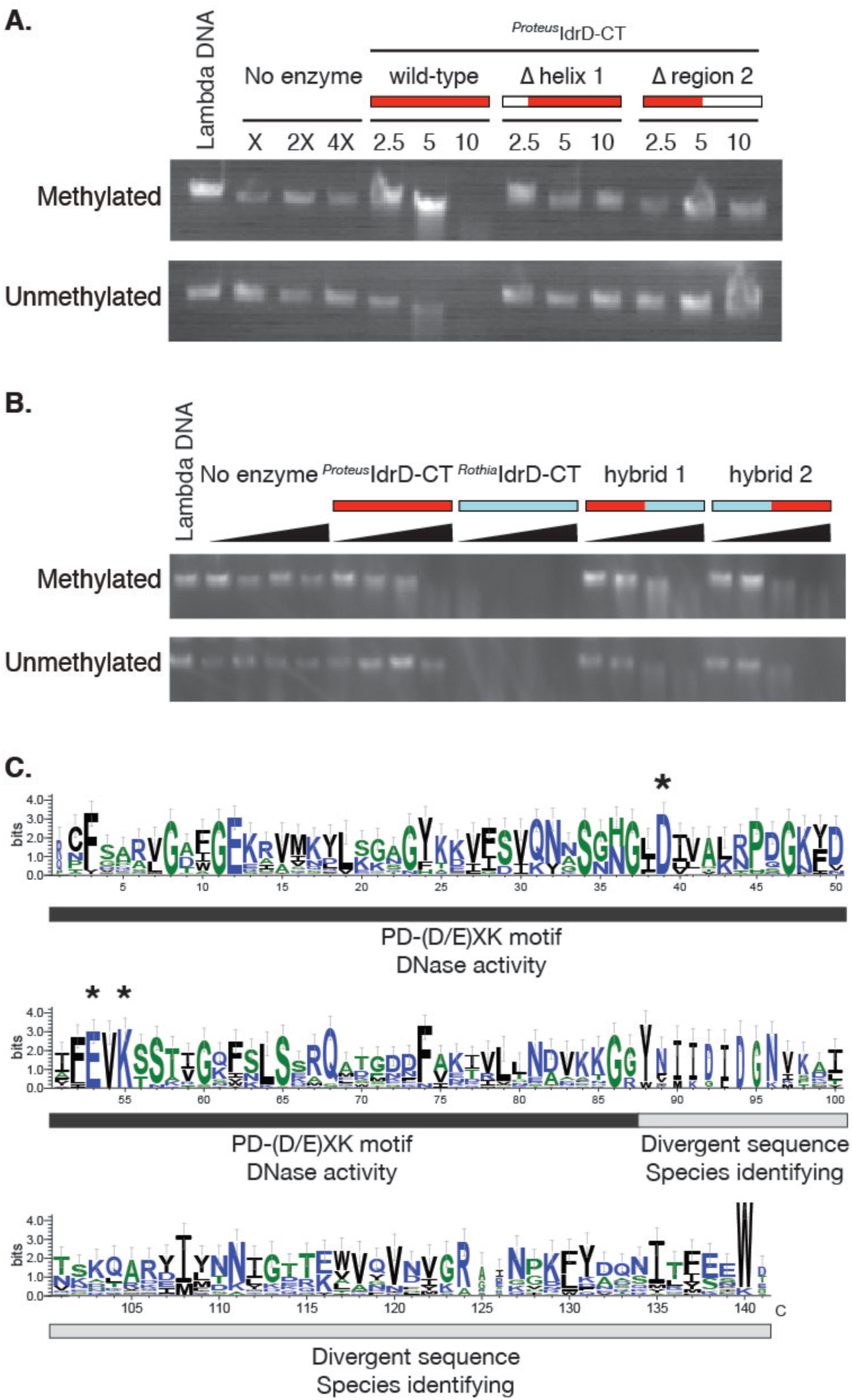
The IdrD-CT proteins are modular, containing independent domains for enzyme activity and species specificity. (A) *In vitro* degradation assay with mutant versions of the *^Proteus^*IdrD-CT protein, none of which lead to DNA degradation. Both domains are essential for DNA degradation. (B) *In vitro* degradation assay with *^Proteus^*IdrD-CT, *^Rothia^*IdrD-CT, and engineered inter-genera hybrid IdrD-CT proteins. All enzymes lead to DNA degradation. (C) WebLogo (*31, 32*) of IdrD-CT and the 23 additional similar proteins with the two domains labeled below.

The variation in the species-identifying domain hinted that the IdrD-CT protein might be modular. If so, then the species-identifying domain should be exchangeable among variants. To test this assertion, we swapped the 3’ region, after the second predicted alpha-helix, between the IdrD-CT proteins of *P. mirabilis* and *R. aeria*. The hybrid IdrD-CT proteins degraded lambda DNA while the negative control did not (Fig. 2B). DNA degradation by these hybrid proteins required more protein than the wild-type enzymes, indicating an attenuation in protein activity (Fig. 2B). As the IdrD-CT proteins are flexible in structural design and can tolerate large domain replacements, we reasoned that one could take advantage of the species-identifying sequences to detect distinct proteins in mixed-species communities.

### IdrD-CT proteins are present in human oral and gut microbiomes

Genes specific to particular strains or species, such as IdrD-CT, could provide a means to identify low-abundance bacteria in deeply-sequenced datasets. Metagenomes are shotgun short-read sequencing of total DNA from a sample that capture a representation of microbial communities, or microbiomes. Bacteria at low abundance in these communities contribute little DNA to the total and often remain hidden. Without sensitive detection methods, the roles and impacts of minority community members remain elusive, including how they defend their niches against more abundant neighbors.

Given the sequence diversity among the IdrD-CT proteins (Fig. 1D), we considered whether these proteins could be used to differentiate between high-and low-abundance bacteria in naturally occurring mixed-species populations. To generate maximum diversity, we focused beyond the IdrD-CT protein and instead on the larger encoding gene. In many of the identified bacteria, including *P. mirabilis* (*10*), the IdrD-CT protein is encoded within the 3’ end of the larger *idrD* gene (Fig. 3A). Before proceeding with the analysis, we deployed an *in silico* test to examine whether seven reference *idrD* gene sequences are sufficiently distinct in nucleotide space to unambiguously identify even short fragments. Reference *idrD* sequences were from *P. mirabilis* BB2000, *R. aeria* C6B, *Cronobacter turicensis* z3032, *Prevetolla jejuni* CD3:33, *Acinetobacter baumannii* XH858, *Pseudomonas fluorescens* F113, and *Xanthomonas citri* pv. *malvacearum* XcmN1003. We generated simulated metagenomes *in silico* from these seven *idrD* seed sequences under different sequencing and error rate scenarios and mapped these onto 127 different combinations of the *idrD* sequences. The mapping revealed no discernible fluctuations in coverage of hits (Fig. S4), supporting that this method allows enough specificity to confidently distinguish between *idrD* sequences originating from different genera.

**Fig. 3.**
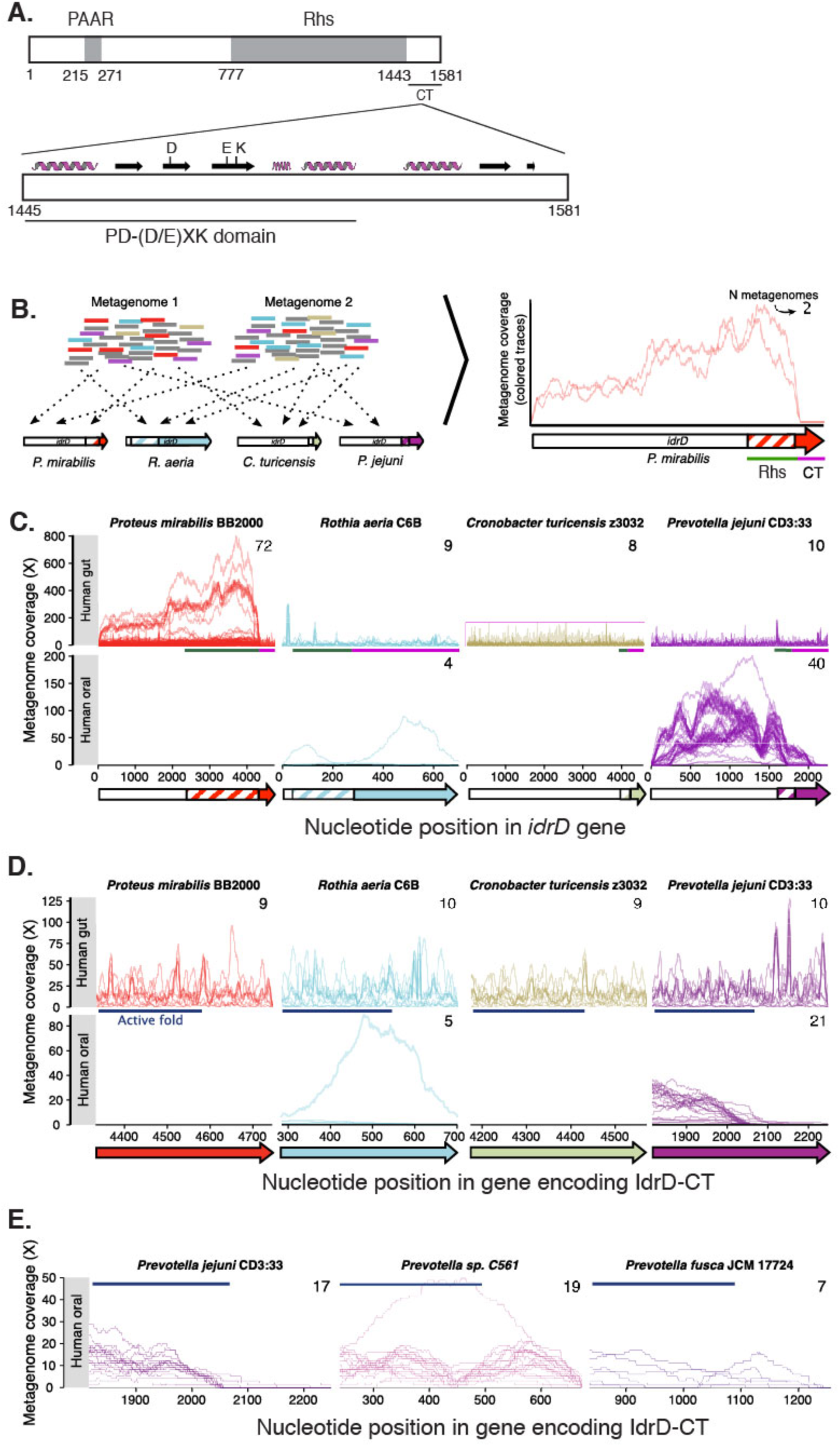
Metagenomics mapping of genes encoding IdrD-CT reveal identifiable differences between bacterial species. (A) At top is a schematic of the IdrD protein, and at bottom is a schematic of the IdrD-CT protein, both drawn to scale. Amino acid numbers are along the bottom, and gray boxes denote conserved PAAR and Rhs domains in the N-terminal region. The predicted secondary structure of IdrD-CT is shown above. (B) Cartoon workflow shows how short reads from metagenomes are mapped against a reference containing the four *idrD* sequences shown here. Based on the mapping results, plotted is each positions’ coverage, i.e., the number of metagenomic reads mapping to that position, for each metagenome (colored, partially transparent lines; each line is a different metagenome). (C) Coverage of each sequence (subpanel columns) by metagenomes originating from the human gut (top row of subpanels) or human oral cavity (bottom row), for metagenomes covering at least half of that *idrD* sequence. Annotated are the positions corresponding to Rhs with dark green bar between the two rows of subpanels and in the striped box on the cartoon depiction of the *idrD* gene. Purple bars mark the positions corresponding to the IdrD-CT proteins. A colored arrow denotes the gene. (D) Higher resolution view of the x-axis to show IdrD-CT coverages by metagenomes covering at least half of that sequence. (E) Shown is coverage for IdrD-CT of three different *Prevotella* species’ *idrD* sequences from metagenomes covering at least half of that sequence.

We next examined metagenomic datasets, capitalizing on the diversity in the nucleotide sequences. We mapped 319 high-quality metagenomes from the human gut or oral microbiomes, onto the *idrD* sequence with the highest sequence similarity from the pool of all seven reference *idrD* sequences (Fig. 3B). This competitive mapping approach maximizes specificity. Plotted on a resultant graph is the number of times a read mapped to a position in *idrD*, termed ‘coverage’ (Fig. 3B). Each line is one metagenome. Smoothly decreasing coverage at the gene ends is an artifact of the mapping process (Supporting Fig. S4C). Coverage otherwise reflects the proportional abundance of a region within the *idrD* gene in a single microbiome sample. We found that many of the sampled human microbiomes harbor the *idrD* gene.

The abundances of each *idrD* sequence generally matched the reported environmental niches of each organism. For example, *Proteus idrD* sequences are abundant in metagenomes from the gut but not for the oral cavity (Fig. 3C). The *idrD* sequences of *A. baumannii* (associated with hospital-acquired infections), *P. fluorescens* (a plant-associated bacterium), and *X. citri* pv. *malvacearum* (a plant pathogen), obtained low coverage across all human samples (Supporting Fig. S5); we removed these organisms from further analysis. The *idrD* sequence of *Prevotella* was also present in oral metagenomes as expected given that the oral microbiota contains numerous *Prevotella* species. Therefore, this mapping reveals abundance and differentiates between genera.

The coverage mapping highlights domains within this gene. The *idrD* gene has two main regions, which is similar to the encoding genes for many polymorphic effectors (*2, 33*). Starting at the 5’ end, most of the gene contains conserved domains, and a subset of the gene (∼ 450 nucleotides at the 3’ end) encodes the effector (Fig. 3A). While metagenomes showed similarly low coverage across the whole gene for *C. turicensis* and *P. jejuni* in gut microbiomes, the two regions of a stereotypical polymorphic effector are apparent for *P. mirabilis*: there is uneven coverage along the length of the gene (Fig. 3C). The Rhs domain (green bars in Fig. 3C) of *P. mirabilis* had higher coverage relative to the rest of the gene (Fig. 3C). Given the Rhs domain’s relatively conserved nucleotide sequences (*15*), reads mapped to the *idrD* Rhs domain likely include all similar Rhs sequences in the metagenome and possibly from genes other than *idrD*.

The 5’ region of the *P. mirabilis idrD* gene had lower coverage relative to the Rhs domain. Similar patterns are present in the graphs for *Prevotella* in oral microbiomes (Fig. 3C). As expected, the 3’ region containing IdrD-CT (purple bars, Fig. 3D) obtained low coverage in gut metagenomes. For *P. mirabilis*, *R. aeria*, and *C. turicensis*, less than 5% of the 319 metagenomes contained sequences for the IdrD-CT proteins. The exception is *P. jejuni* in which ∼ 10% of microbiomes indicated the presence of genes encoding an IdrD-CT protein, suggesting that these proteins occur at low frequency in humans. Therefore, we conclude that multiple variants of *idrD*-like genes are present in microbiome samples, yet only a subset of these genes encode an IdrD-CT protein.

The coverage pattern for the *P. jejuni* IdrD-CT fell approximately halfway through the encoding sequence, leading us to consider whether this pattern reflects true sequence variation among *Prevotella* species. The region containing the PD-(D/E)XK domain recruited reads abundantly from oral microbiomes, while the remainder of the sequence obtained comparably fewer reads (Fig. 3D). Given that *P. jejuni* was isolated from the human gut, we reasoned that the 3’ region sequence within the gene encoding IdrD-CT could differ from that of *Prevotella* species inhabiting the mouth. Therefore, we undertook an additional mapping analysis on the same oral metagenomes except using the nucleotide sequences of IdrD-CT from *P. sp.* C561 and *P. fusca* JCM 17724; both are isolates from the human oral cavity. *P. sp.* and *P. fusca* sequences received relatively even coverage across the entire sequence (Fig. 3E). The *P. jejun*i sequence, which was added into the mapping analysis, again recruited little coverage in the 3’ region of IdrD-CT (Fig. 3E). The 3’ region of IdrD-CT thus reflects species diversity within the *Prevotella* genus.

## Discussion

Effector families point to competitive strategies likely optimized for a particular niche or community composition. Earlier metagenomic analysis on the abundance of cell-contact dependent toxic proteins, often originating from a single species, suggested that these effectors help establish dominance in a community and predicted niche-specific specialization of these proteins (*8, 9*). Emerging is the possibility that genes in low-abundance and specific for species (or strains) could open a window for differentiating among bacterial populations within a complex community. To further address this hypothesis, the abundance of low-abundance species should be evaluated by incorporating greater resolution of species-identifying protein regions.

To this end, we have described IdrD-CT, a *P. mirabilis* self-recognition protein that acts on adjacent cells through direct contact (*10*). IdrD-CT is the founding member for a family of DNases; these proteins are modular and contain an interchangeable species-identifying domain. We propose to term this family, “Idr-PDE-DNase.” Through a combination of genetics and molecular biology along with biochemical and phylogenetic interrogation, we showed that IdrD-CT from *P. mirabilis* and *R. aeria* degrade DNA regardless of methylation state (Fig. 1). Idr-PDE-DNase proteins are found in evolutionarily distant bacteria and together form a distinct family of the PD-(D/E)XK nuclease superfamily. Finding and characterizing additional families is crucial for optimizing methods to predict function from sequence for this superfamily.

Further, we demonstrated that metagenomic analysis can provide fine functional resolution of the Idr-PDE-DNase proteins in human samples, even when sequences are present at low frequency in datasets. For example, the 3’ region of the IdrD-CT nucleotide sequence distinguished gut and oral *Prevotella* species (Fig. 3E), reflecting diversity in bacterial populations naturally occurring in human guts and oral cavities. Flexibility in sequence space for this 3’ region might allow each Idr-PDE-DNase protein to become distinct among recently diverged groups such as species.

How gene clusters encoding strain-specific proteins (e.g., toxins or effectors) evolve is an intriguing question in multiple contexts. Two-pair toxic protein systems are analogous to colonization and addiction proteins or mating compatibility such as in *Wolbachia* (*34*) and in *Caenorhabditis elegans* (*35*). One current hypothesis is that speciation occurs on the gene cluster of the effector protein (toxin) and its immunity protein (anti-toxin), for example, through horizontal gene transfer, as these two-gene pairs are often in variable hotspots on the genome (*2, 3, 7, 15*). Yet for these Idr-PDE-DNase proteins, a pair of domains within a protein follow such a tandem pattern, raising the possibility that an evolutionary arms race, internal to the protein, may also drive speciation. Frequent horizontal gene transfer and/or shifting selection pressures could also explain the sequence variability within the Idr-PDE-DNase proteins.

Of the two essential domains of the Idr-PDE-DNase proteins, one has enzyme activity and is conserved, while the other is variable and contains information specific to genus and species. A need to maintain enzymatic activity (*22, 36–40*) might restrict the conserved enzymatic domain from changing. By contrast, the C-terminal half of Idr-PDE-DNase proteins vary in amino acid sequences (Fig. 2C). Similar DNA-targeting proteins act as multimeric complexes, and similar effector proteins can bind allele-specific immunity proteins (*2, 3, 7, 41*). The C-terminus of Idr-PDE-DNase proteins might contribute to protein-binding and/or DNA-protein interactions, especially as the predicted secondary structures are similar among these proteins. The functional exchange of the C-terminal region between two Idr-PDE-DNase (from *Proteus* and *Rothia*) proteins are consistent with this hypothesis (Fig. 2B).

Unlike chimeric nucleases such as homing nucleases, zinc finger nucleases, TALENS, and CRISPR/Cas9 (reviewed in (*42, 43*)), the DNA recognition motif for the Idr-PDE-DNase proteins is unknown. Moreover, the non-nuclease domain does not seem to contain a typical DNA-binding motif. Structural studies are needed to discern how Idr-PDE-DNase proteins differ in interactions with DNA and protein partners. The modularity in function and flexibility in amino acid sequences open the Idr-PDE-DNase proteins to possible engineering and for the development of clinical tools.

Further, capitalizing on fine-detail differences as species-identifying markers allows one to pull out rare species, and perhaps strains, in naturally occurring communities. For example, the sequence differences in the variable region of IdrD-CT acted as tokens for identifying distinct strains or species in metagenomic datasets. Molecular identification of sequence variants with known functional consequences also informs physiologically relevant population boundaries when resolved in large sequence datasets. Combining metagenomic analysis with molecular characterization of strain-identifying factors opens a window into understanding how interactions among resident microbes might contribute to behaviors within a host.

## Acknowledgments

We thank Colleen Cavanaugh and Philippe Cluzel for comments on this manuscript and Alecia Septer for preliminary experiments. **Funding:** This research was funded by the David and Lucile Packard Foundation, the George W. Merck Fund, and Harvard University. Additional support was provided to D.R.U by Harvard University’s Department of Organismic and Evolutionary Biology program and the National Science Foundation Graduate Research Fellowship Program under Grant No. DGE1745303 to D.R.U.

## Author contributions

D.S. performed all experiments. D.R.U. performed all computational analysis. D.S. and K.A.G. developed this project. D.S. performed all experiments. D.U. performed all computational analysis. D.S. and D.R.U. led the experimental and computational design, respectively, with contributions by K.A.G. and each other. All three authors wrote and edited the manuscript.

## Data and materials availability

Strains and primary data are available upon request. Computational methods are available on www.gibbslab.org. Original gels and sequence accession data are provided in the supplementary data.

## Materials and Methods

### Bacterial strains and media

Overnight cultures of all strains were grown aerobically at 37°C in LB broth. *P. mirabilis* strains were maintained on low swarming (LSW^-^) agar (*10*) and allowed to swarm on CM55 agar (blood agar base agar (Oxoid, Basing-stoke, England). *E. coli* strains were grown on 1.5% LB agar. For plasmid maintenance, growth media had 35g/ml kanamycin. All strains are listed in Table S1. Oligos used in this study are listed in Table S2.

### Strain construction

The P*_idrA_*-*idrD* expression plasmid was constructed by using polymerase chain reactions (PCR) to amplify the last 416 base pairs (bp) of the *idrD* gene from BB2000 using primers AS174 and AS175 and cloning it into the SacI and AgeI sites of pAS1034, resulting in plasmid pAS1054. The inducible anhydrotetracycline promoter (Ptet) (*44*) was introduced into the *idrD* expression plasmids by generating a gBlock (gDS0005) of the promoter region with 29 bp overhangs for the plasmid, and using SLiCE (*45*) to recombine into pAS1054, resulting in the plasmid pDS0002 (*idrD*).

A C-terminal FLAG tag (GACTACAAGGACGACGATGACAAG) was added to *idrD* by using SLiCE (*45*) to recombine the gBlock gDS0023 (FLAG tag with 49 bp overhang of *idrD* and 52 bp overhang of pDS0002) into pDS0002, resulting in the plasmid pDS0034. The FLAG-tagged *idrD* active site mutants were generated by replacing *idrD-FLAG* in pDS0034 with the mutant sequences which are encoded in gBlocks gDS0025-28, resulting in plasmids pDS0048 (D39A), pDS0049 (E53A), pDS0050 (K55A), and pDS0051 (triple mutant). The untagged versions of *idrD* were constructed by PCR amplifying the mutant *idrD* sequences from pDS0048-51 with oDS0137 and oDS0159 to remove FLAG tag and performing a restriction digest with SacI and AgeI to insert into pDS0034 (pDS0058-61).

Primers oDS0161 and oDS0162 were used to amplify *gfpmut2* from pids_BB_*-idsE*-GFP. This PCR fragment was inserted into pDS0034 through restriction digest with SacI and AgeI to generate pDS0062, which is a *gfp* expression plasmid under the control of Ptet. The plasmid encoding IdsE-GFP was constructed by single overlap extension of *idsE*, the flexible linker (GSAGSAAGSGEF, (*46*), *gfpmut2* (*47*), and *idsF*, followed by restriction digest cloning into the AgeI (in *idsE*) and KpnI (in *idsF*) unique sites in the plasmid pids_BB_.

All plasmids were confirmed by Sanger sequencing (Genewiz). Plasmids with conjugative transfer elements, including all IdrD-CT expression plasmids, were moved into the *E. coli* conjugative strain S17, which were then mated with recipient *P. mirabilis* strains. The presence of plasmids was confirmed in recipient strains by PCR using plasmid-specific primers.

### Liquid viability assay

Overnight cultures of *E. coli* MG1655 carrying each expression plasmid were normalized to an optical density at 595 nm (OD_595_) of 1.2 µL of normalized cultures was added to 198 µL of LB broth supplemented with kanamycin and 10 nM anhydrotetracycline and grown at 37°C for 16 hours in a 96 well-plate. OD_595_ readings were taken every half hour.

### Microscopy

We performed microscopy on *P. mirabilis* strain BB2000 carrying either vector pBBR1-NheI (*48*) or pDS0002 (producing IdrD-CT) and on *E. coli* carrying either pBBR1-NheI (*48*), pDS0002, or pDS0048 (producing IdrD-CT_D39A_). *P. mirabilis* cells were normalized to OD 0.1 after overnight growth in LB supplemented with kanamycin. Cells were inoculated onto CM55 swarm pads containing 10 µg/mL DAPI and 10 nM anhydrotetracycline and grown in humidified chambers at 37°C. At five and six hours after growth. From overnight cultures, *E. coli* cells were grown in LB plus kanamycin until mid-logarithmic phase and then mounted directly onto glass slides. Glass coverslips were sealed with nail polish. For all microscopy, we captured phase contrast and DAPI (150 ms exposure) images using a Leica DM5500B microscope (Leica Microsystems, Buffalo Grove IL) and CoolSnap HQ CCD camera (Photometrics, Tucson AZ) cooled to -20°C. MetaMorph version 7.8.0.0 (Molecular Devices, Sunnyvale CA) was used for image acquisition.

### *In vitro* DNase assay

IdrD and IdrD-CT_D39A_ with a C-terminal FLAG epitope tag were produced using the New England Biolabs PURExpress *In Vitro* Protein Synthesis Kit. Template DNA was amplified from pDS0034 or pDS0048 using primers with overhangs to add the required elements specified by the PURExpress kit. Reactions were performed with 250 ng of template DNA (no template DNA added to negative control reaction) and incubated at 37°C for two hours. Protein amount was determined using an anti-FLAG western blot with a known gradient of FLAG-BAP (2.5, 5, 10, and 20 ng). Synthesized protein (2.5, 5, and 10 ng) was added to 0.5 µg of lambda DNA (methylated and unmethylated), 5 µL of New England Biolabs Buffer 3.1, and up to a final volume of 25 µL. For plasmid DNase assays, 10 ng of synthesized protein was added to 250 ng of circular or linear plasmid DNA (pids_BB_). This reaction was incubated for one hour at 37°C, then Proteinase K (New England Biolabs) was added and incubated for 15 minutes at 37°C. Reaction was then run on a 1% agarose gel for analysis.

### Western blotting

Samples were run on a 12% Tris-Tricine polyacrylamide gel, transferred to a nitrocellulose membrane, probed with rabbit anti-FLAG (1:4,000; Sigma-Aldrich, St. Louis, MO), then goat anti-rabbit conjugated to horseradish peroxidase (HRP) (1:5,000; KPL, Inc., Gaithersburg, MD), and finally developed using Immun-Star HRP substrate kit (Bio-Rad Laboratories, Hercules, CA). Blots were visualized using a Chemidoc (Bio-Rad Laboratories, Hercules, CA) and TIFF files were used for analysis on Fiji (ImageJ, Madison, WI).

### Identification of proteins with similarity to IdrD-CT

The amino acid sequence of *^Proteus^*IdrD-CT was used to search publicly available protein databases using phmmer (*24, 25*). Additionally, the nucleotide sequence of *^Proteus^*IdrD-CT was used to search against nucleotide sequence databases using tblastn (*26, 27*). By hand, the amino acid sequences obtained from these searches were checked for the presence of the predicted active site residues and the absence of truncations compared to the *^Proteus^*IdrD-CT. The full C-terminal domain was further determined by looking for an Rhs or VENN motif-containing domain, which marks the start of the C-terminal domain (*2, 7, 15*). Alignments were of IdrD-CT and similar IdrD-CTs performed using MUSCLE followed by Ali2D on MPI Bioinformatics toolkit (*23*)

### Phylogenetic reconstruction of *idrD* and species relationships

The amino acid sequences for 25 *idrD* genes identified as described above were obtained and aligned with muscle (*49*) before removing positions with less than 70% occupancy with trimAl (*50*). This alignment was passed to MrBayes v3.2.6 (*51*) to reconstruct the tree using the WAG substitution model (*52*) and gamma model of rate heterogeneity. Each run has 20 million generations sampled every 1,000 generations by four coupled chains heated at the default temperature of 0.2. Four independent runs were checked for convergence and then combined into a 50% majority rule tree after burning the initial 40% of the sampled trees. The species tree was reconstructed using identical methods but with full-length 16S ribosomal RNA sequences obtained from the various genomes, choosing arbitrarily if multiple 16S rRNA copies were found in a genome. Specifically, alignment and MrBayes parameters were identical except for changing the model to GTR (*53*). Maximum-likelihood trees were also generated with RAxML (*54*) using the appropriate WAG or GTR + gamma models and produced identical topologies, albeit with less support (data not shown).

### Metagenome selection and processing

A fully-reproducible workflow explaining and documenting commands and scripts used to perform the metagenome abundance is available at www.gibbslab.org on the Resources page. Full-length *idrD* nucleotide sequences were obtained from *Acinetobacter baumannii* XH858, *Cronobacter turicensis* z3032, *Prevotella jejuni* CD3:33, *Proteus mirabilis* BB2000, *Pseudomonas fluorescens* F113, *Rothia aeria* C6B, and *Xanthomonas citri* pv. *malvacearum* XcmN1003. Publicly available metagenomes likely to represent populations with these genes were identified using the Search SRA portal (www.searchsra.org) that employs the PARTIE algorithm (*55*). Briefly, PARTIE uses bowtie2 (*56*) to search a 100,000-read random subset from each of ∼110,000 published metagenomes. From this, 3,801 candidate metagenomes were identified that contributed non-zero coverage to at least one *idrD* sequence, from which we curated a list of 1,189 metagenomes by removing genome assemblies, transcriptomes, non-random library preparation methods, etc. 1,188 of these metagenomes were downloaded from available on NCBI’s Short Read Archive (fastq-dump --split-3) for deeper analysis (1 metagenome could not be downloaded). A single bowtie2 database was created with all seven *idrD* nucleotide sequences, onto which bowtie2 mapped metagenomes using default parameters (--sensitive; (*56*)).

Anvi’o, an analysis and visualization platform for ‘omics data, managed the resultant data and subsequent abundance (*57*). With Anvi’o, a contigs database was generated (anvi-gen-contigs-database command) from the seven *idrD* sequences and profiled with the merged results of the bowtie2 mapping (anvi-profile and anvi-merge commands, respectively). Per-nucleotide coverages and variability (i.e., single-nucleotide polymorphism (SNP) counts) were exported for all sequences with the Anvi’o anvi-get-split-coverages and anvi-gen-variability-profile commands, respectively.

To investigate the distribution of *idrD* coverage among different *Prevotella* in the human oral cavity, *idrD* sequences from the three additional *Prevotella* spp. (*P.* sp. C561, *P. fusca* JCM 17724, *P. denticola* NCTC 13067) along with the *P. jejuni* CD3:33 *idrD* sequence were mapped separately, using the same methods, mapping reads from 202 human oral cavity metagenomes.

### *In silico* metagenome simulation

Short-read metagenome sequences were simulated from all seven full-length *idrD* reference nucleotide sequences with the ART simulator (Huang et al., 2012). We modeled two different scenarios; one based on using an Illumina HiSeq 2000 with 100x the normal error rate to sequence a metagenome to a depth of 30x with 25bp single-end reads short reads (Supplemental Figure S3). The other scenario (Supplemental Figure S3C) was modeled with identical parameters except with the normal HS2000 error rate and paired-end 100bp reads. Each of the 127 different subset combinations of the seven full-length *idrD* reference sequences was used to create reference databases containing a different subset of available reference sequences, to which each artificially generated metagenome was mapped. Supplemental Figure S3 then shows the coverage for each sequence (columns) for the different combination categories (rows).

### Metagenome categorization

Metagenomes were binned into categories based on the provided “ScientificName” annotation. Metagenomes from the Human Microbiome Project (HMP) (*58, 59*) were listed as “human metagenome”; these were disaggregated into “HMP oral metagenome” and “HMP gut metagenome” based on the sampled site listed in the “Analyte_Type” column. From the 1,188 metagenomes, we focused on only the 324 human oral or human gut metagenomes, defining human oral as metagenomes labelled “human oral metagenome”, “HMP oral metagenome”, or “Non-HMP oral metagenome” (n = 42) and human gut as metagenomes labelled “human gut metagenome” or “human metagenome” (n = 277). Metagenomes annotated as “human metagenome” (n = 47) were categorized as human gut since though they originated from a variety of human body sites, the only three passing the filtration criteria (see next section) were from the human gut. Metagenomes annotated as “oral metagenome” (n = 112) were excluded as extremely low (single digit), noisy coverages were recovered. In addition, five metagenomes were specifically discarded: SRR628272 was removed from all datasets as the original FASTQ had extremely low quality scores; SRR1779144 came from a diseased infant (https://www.ncbi.nlm.nih.gov/sra/?term=SRR1779144) and skewed the y-axis with an extremely high *C. turicensis* coverage (>1,000x); SRR2047620 came from a pediatric stem-cell treatment dataset with high antibiotic loads (https://www.ncbi.nlm.nih.gov/sra/?term=SRR2047620) with similarly extreme *C. turicensis* coverage; SRR1781983 came from a subgingival plaque of a patient with periodontitis (https://www.ncbi.nlm.nih.gov/biosample/SAMN03287617) and had extremely high *R. aeria* coverage that skewed the y-axis (>500x); and SRR1038387 came from an infant with necrotizing enterocolitis (https://www.ncbi.nlm.nih.gov/sra/?term=SRR1038387) with extremely high *P. mirabilis* coverage that skewed the y-axis (>2,000x). 1,183 metagenomes remained after discarding these metagenomes, of which 319 were human oral or human gut metagenomes. Moving forward, we focused exclusively on *P. mirabilis, R. aeria*, *C. turicensis*, and *P. jejuni* as the other taxa’s sequences had little to no coverage from the human metagenomes of interest.

### Mapping filtration criteria

To minimize noisy coverage originating from metagenomes relevant for one organism but not another, we employed a filtering strategy that, for each *idrD* sequence, considered only metagenomes from which at least half of the nucleotides received coverage. For Figures 4C and 4D, the filtering criterion was applied after subsetting to the C-terminal domain (CTD); that is, the filtration was applied based only on the CTD. The number of metagenomes passing the criterion is displayed in the top right corner of each subpanel in Figure 4. For each metagenome passing the filtration step, each nucleotides’ coverage is plotted as a partially-transparent line; thus, metagenomes with similar coverage trends overlap and appear darker. Metagenome coverages inherently decrease smoothly towards the 3’ and 5’ ends of the gene because, moving towards the ends, the probability decreases that a short read will overlap enough of the reference sequence to be mapped there.

---

**Figure S1:**
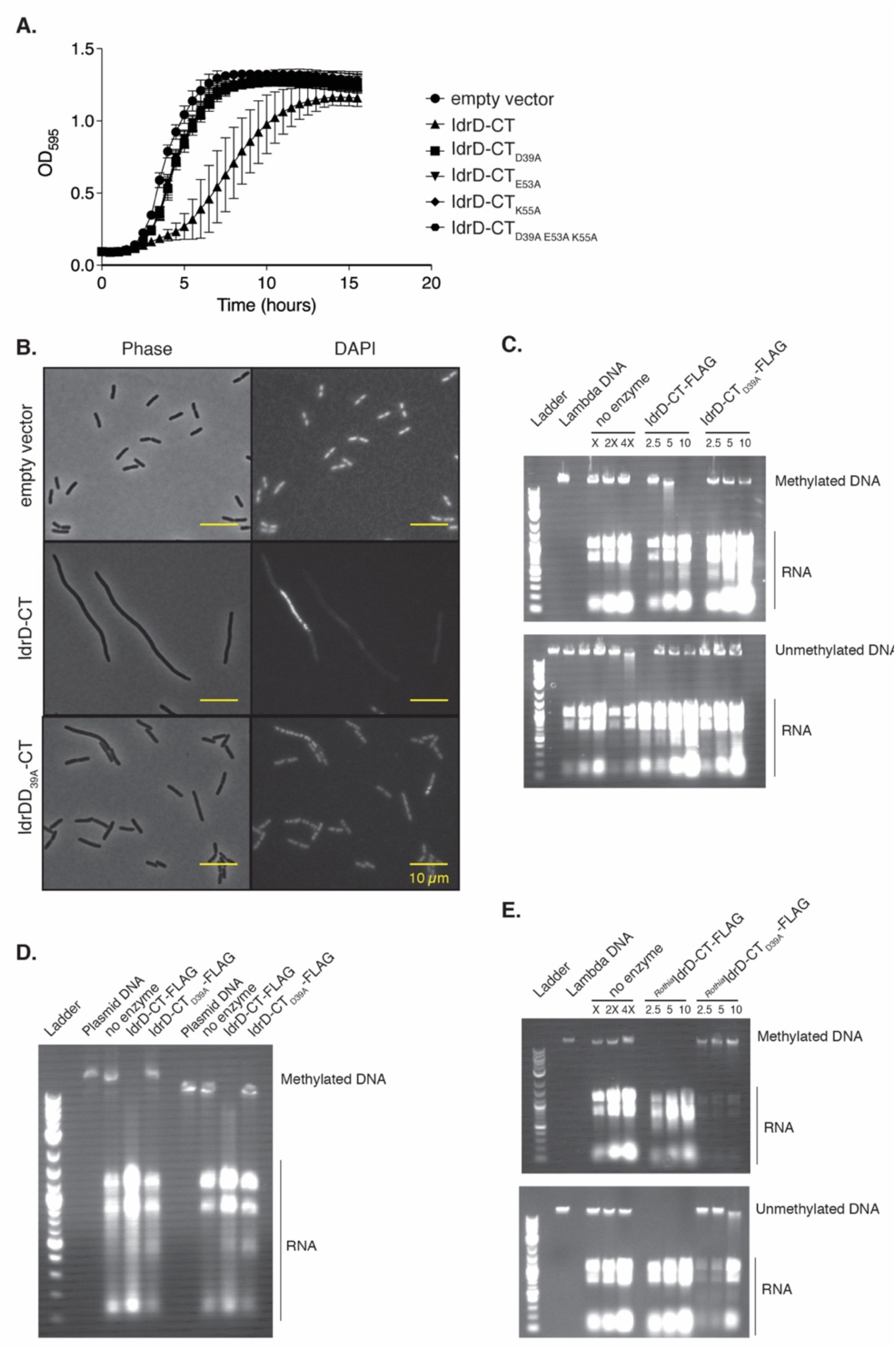
***P. mirabilis* IdrD-CT is an endonuclease in the PD-(D/E)XK superfamily.** (A) Quantification of viable cells after overexpression of IdrD-CT and active site mutants in liquid-grown *E. coli* MG1655. Optical density at 595 nm over the course of eight hours are plotted on a log_10_ scale. (B) The micrographs show *E. coli* cells isolated from mid-logarithmic growth in LB plus kanamycin. Top, the parent plasmid as a negative control. Middle, wild-type IdrD-CT of *P. mirabilis* strain BB2000, produced on the plasmid. Bottom, the null mutant IdrD-CT_D39A_ of *P.* mirabilis strain BB2000, produced on the plasmid. Left, Phase. Right, fluorescence of DAPI-stained DNA. (C and D) Full agarose gels of DNase assays with (C) lambda DNA as substrates, with NEB 2-log DNA ladder and (D) plasmid DNA. Increasing amounts (2.5, 5 and 10 ng) of IdrD-CT-FLAG (red box) and IdrD-CT_D39A_-FLAG (pink with black star) were added to methylated and unmethylated lambda DNA (48,502 bp). The negative control with no protein produced is depicted with a white box. DNA is degraded in the samples with IdrD-CT-FLAG. Bands running below 1kB are presumed to be rRNA and tRNA from PURExpress reaction. (E) *In vitro* DNA assays with *Rothia*-produced IdrD-CT-FLAG and IdrD-CT_D39A_-FLAG.

**Figure S2:**
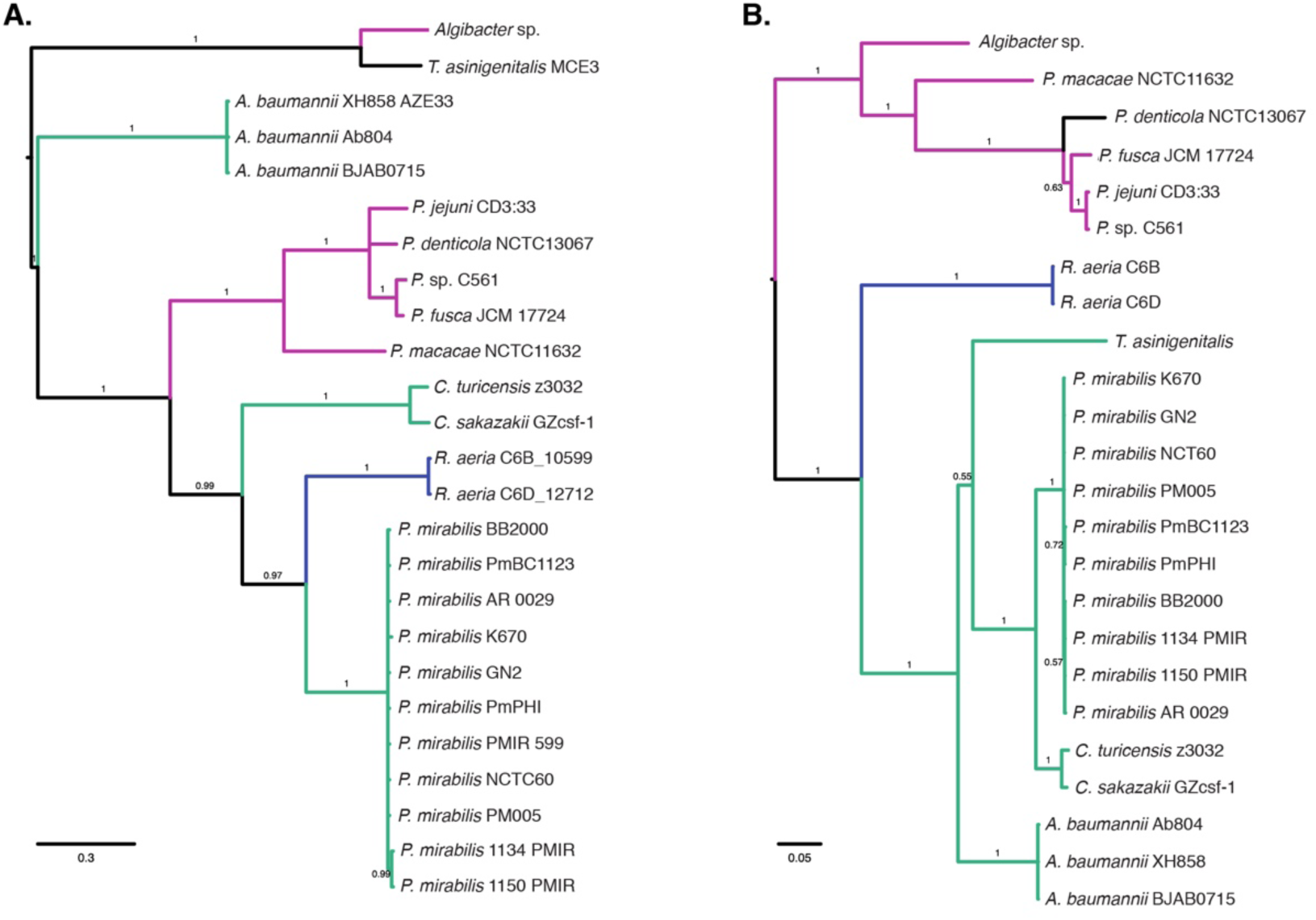
Proteins with similarity to IdrD-CT are found in evolutionarily distant bacteria. (A) Phylogeny of proteins identified as similar to IdrD-CT. The amino acids of these sequences are aligned in Figure 1. (B) Species tree of taxa containing proteins with similarity to IdrD-CT. 16S rRNA sequences were obtained from published genomes of the taxa represented in B. In both A and B, branches are colored by phyla (pink: Bacteriodetes, teal: Proteobacteria, blue: Actinobacteria). Numbers adjacent to each branch report posterior probability. Scale bars show expected substitutions per site.

**Figure S3:**
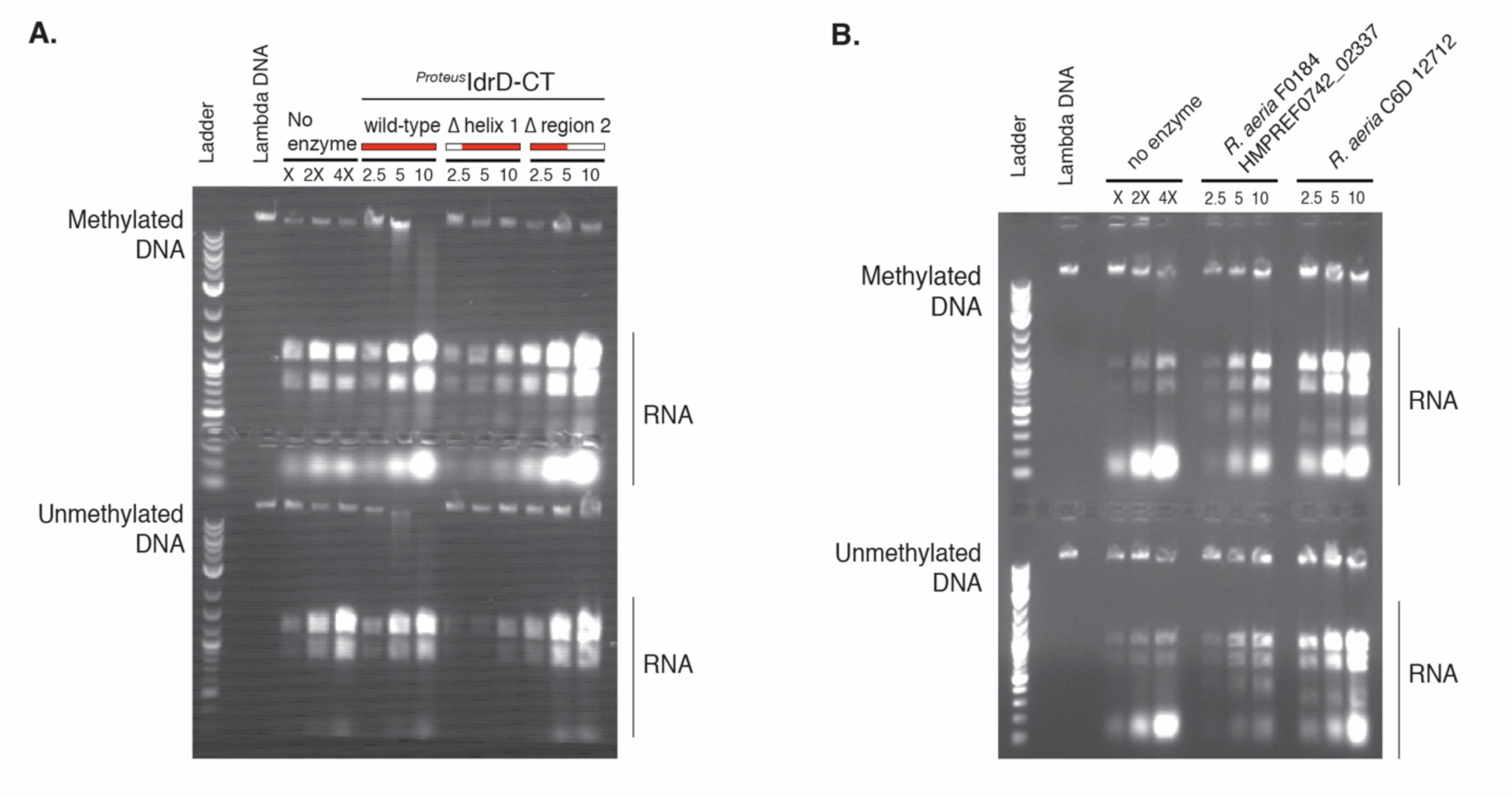
D**N**A **degradation requires full-length of IdrD-CT.** Increasing amounts (2.5, 5 and 10 ng) of protein were added to methylated and unmethylated lambda DNA (48,502 bp) and run on an agarose gel with NEB 2-log DNA ladder. Bands running below 1kB are presumed to be rRNA and tRNA from PURExpress reaction. The negative control with no produced protein is depicted with a white box. Pure lambda DNA is labeled as such. (A) N-terminal truncations (IdrD-CT-FLAG 16-137) and C-terminal truncations (IdrD-CT-FLAG 1-79) of *P. mirabilis* IdrD-CT, and (B) IdrD-CTs from *Rothia* spp. with N- and C-terminal truncations. Deleted regions are depicted with a white box.

**Figure S4:**
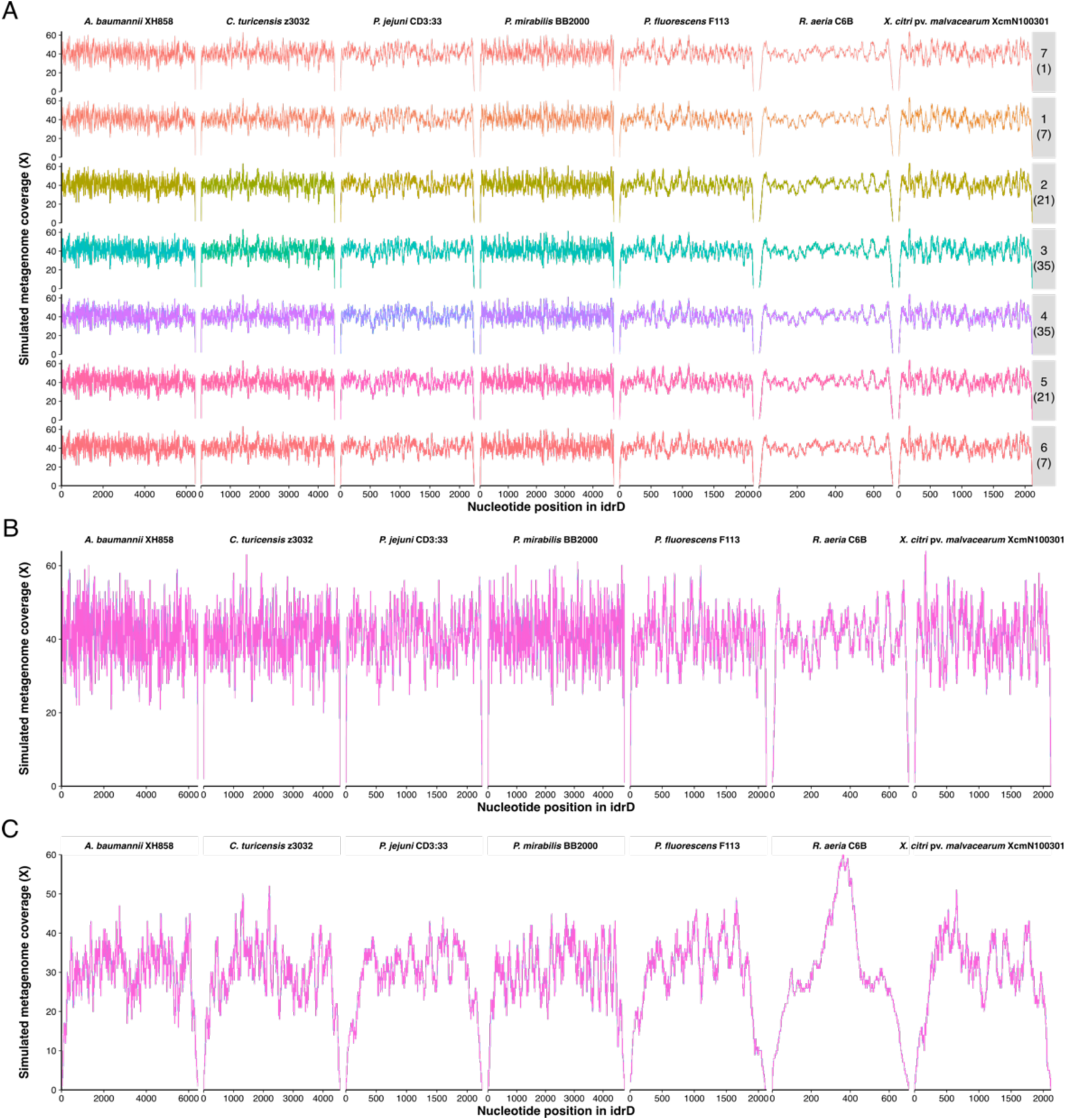
*In silico* metagenome simulation reveals that seven *idrD* DNA sequences are distinguishable in datasets of short-read DNA sequences. (A) Short reads were modeled with art_illumina to model a metagenome sequenced on an Illumina HiSeq 2000 to a depth of 30x with 25bp single-end reads but with 100x the normal error rate. The individual plots show the coverage attained by that *idrD* sequence (columns) for a variety of reference combinations (rows). Each row represents a different number of *idrD* sequences in the reference set (top number in grey box), with all combinations choosing that number of sequences represented (bottom number in parentheses) for a total of 127 different combinations. For example, the 3^rd^ row from the top reports all 21 mapping analyses onto the 21 different combinations of two-sequence reference databases. Each trace shows the coverage of an individual mapping analysis combination and is colored with a rainbow palette. (B) Same data as in A but all columns are collapsed to show that there is no difference by mapping combination. (C) Same analysis as in B but using a metagenome simulated based on an Illumina HiSeq 2000 sequencer with paired-end 100bp reads and normal error rate.

**Figure S5.**
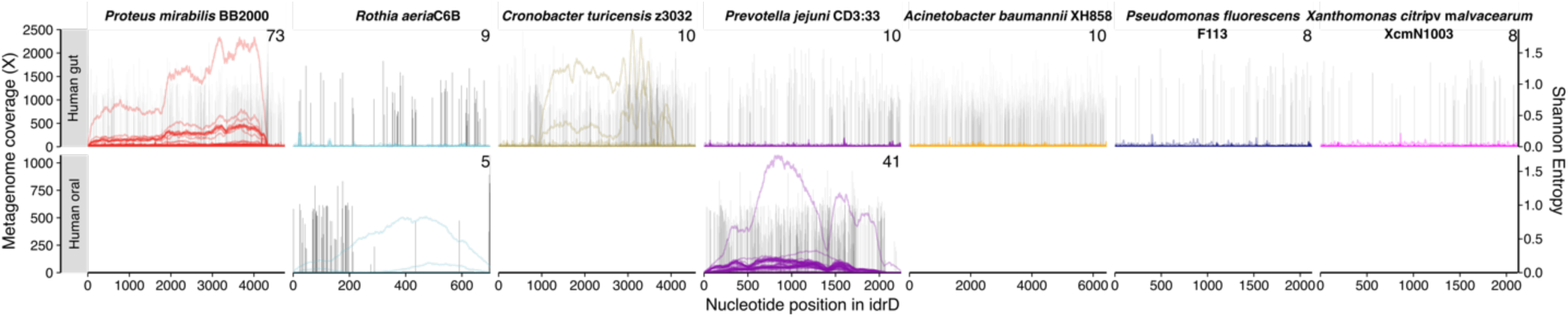
The *idrD* sequences from *Acinetobacter baumannii*, *Pseudomonas fluorescens,* and *Xanthomonas citri* pv. *malvacearum idrD* sequences are not abundant in human oral or gut metagenomes. Coverage of each sequence (subpanel columns) by metagenomes originating from the human gut (top row of subpanels) or human oral cavity (bottom row), for metagenomes covering at least half of that *idrD* sequence. The number of metagenomes passing this filter is shown in the top right corner of each subpanel. No samples have not been removed from this analysis.

**Table S1.** **Strains and plasmids.**

| Strain | Notes | Strain name | Source |
| --- | --- | --- | --- |
| *P. mirabilis*<br>BB2000 *idrD* * +<br>pDS0062 | *idrD::Tn-Cm(R)* producing GFPmut2 under the control of an anhydrotetracycline-inducible promoter (pBBR1 origin, Kan (R)) | DS349 | This study |
| *P. mirabilis*<br>BB2000 *idrD* * +<br>pDS0002 | *idrD::Tn-Cm(R)* producing IdrD-CT under the control of an anhydrotetracycline -inducible promoter (pBBR1 origin, Kan (R)) | DS104 | This study |
| *P. mirabilis*<br>BB2000 *idrD* * +<br>pDS0058 | *idrD::Tn-Cm(R)* producing the D39A mutant variant of IdrD-CT under the control of an anhydrotetracycline -inducible promoter (pBBR1 origin, Kan (R)) | DS344 | This study |
| *P. mirabilis*<br>BB2000 *idrD* * +<br>pDS0059 | *idrD::Tn-Cm(R)* producing the E53A mutant variant of IdrD-CT under the control of an anhydrotetracycline -inducible promoter (pBBR1 origin, Kan (R)) | DS345 | This study |
| *P. mirabilis*<br>BB2000 *idrD* * +<br>pDS0060 | *idrD::Tn-Cm(R)* producing the K55A mutant variant of IdrD-CT under the control of an anhydrotetracycline -inducible promoter (pBBR1 origin, Kan (R)) | DS346 | This study |
| *P. mirabilis*<br>BB2000 *idrD* * +<br>pDS0061 | *idrD::Tn-Cm(R)* producing the triple mutant variant (D39A, E53A, K55A) of IdrD-CT under the control of an anhydrotetracycline - inducible promoter (pBBR1 origin, Kan (R)) | DS347 | This study |
| *E. coli* MG1655<br>+ pBBR1-NheI | MG1655 carrying an empty plasmid (pBBR1 origin, Kan (R)) | DS68 | This study |
| *E. coli* MG1655<br>+ pDS0002 | MG1655 producing IdrD-CT under the control of an anhydrotetracycline-inducible promoter (pBBR1 origin, Kan (R)) | DS151 | This study |
| *E. coli* MG1655<br>+ pDS0058 | MG1655 producing the D39A mutant variant of IdrD-CT under the control of an anhydrotetracycline-inducible promoter (pBBR1 origin, Kan (R)) | DS336 | This study |
| *E. coli* MG1655<br>+ pDS0059 | MG1655 producing the E53A mutant variant of IdrD-CT under the control of an anhydrotetracycline-inducible promoter (pBBR1 origin, Kan (R)) | DS337 | This study |
| *E. coli* MG1655<br>+ pDS0060 | MG1655 producing the K55A mutant variant of IdrD-CT under the control of an anhydrotetracycline-inducible promoter (pBBR1 origin, Kan (R)) | DS338 | This study |
| *E. coli* MG1655<br>+ pDS0061 | MG1655 producing the triple mutant variant (D39A, E53A, K55A) of IdrD-CT under the control of an anhydrotetracycline-inducible promoter (pBBR1 origin, Kan (R)) | DS339 | This study |
| *P. mirabilis*<br>BB2000 +<br>pBBR1-NheI | Wild-type *P.mirabilis* carrying an empty<br>vector (pBBR1 origin, Kan (R)) |  | (48, 60) |
| *P. mirabilis*<br>BB2000 +<br>pDS0002 | *idrD::Tn-Cm(R)* producing IdrD-CT under<br>the control of an anhydrotetracycline -<br>inducible promoter (pBBR1 origin, Kan (R)) |  | This study |
| OneShot<br>Omnimax 2 T1R<br>Competent Cells | *E. coli* strain for cloning |  | Thermo<br>Fisher<br>Scientific,<br>Waltham,<br>MA. |
| S17λpir | *E. coli* mating strain to introduce plasmids<br>into *P. mirabilis* |  | (61) |

**Table S2.**
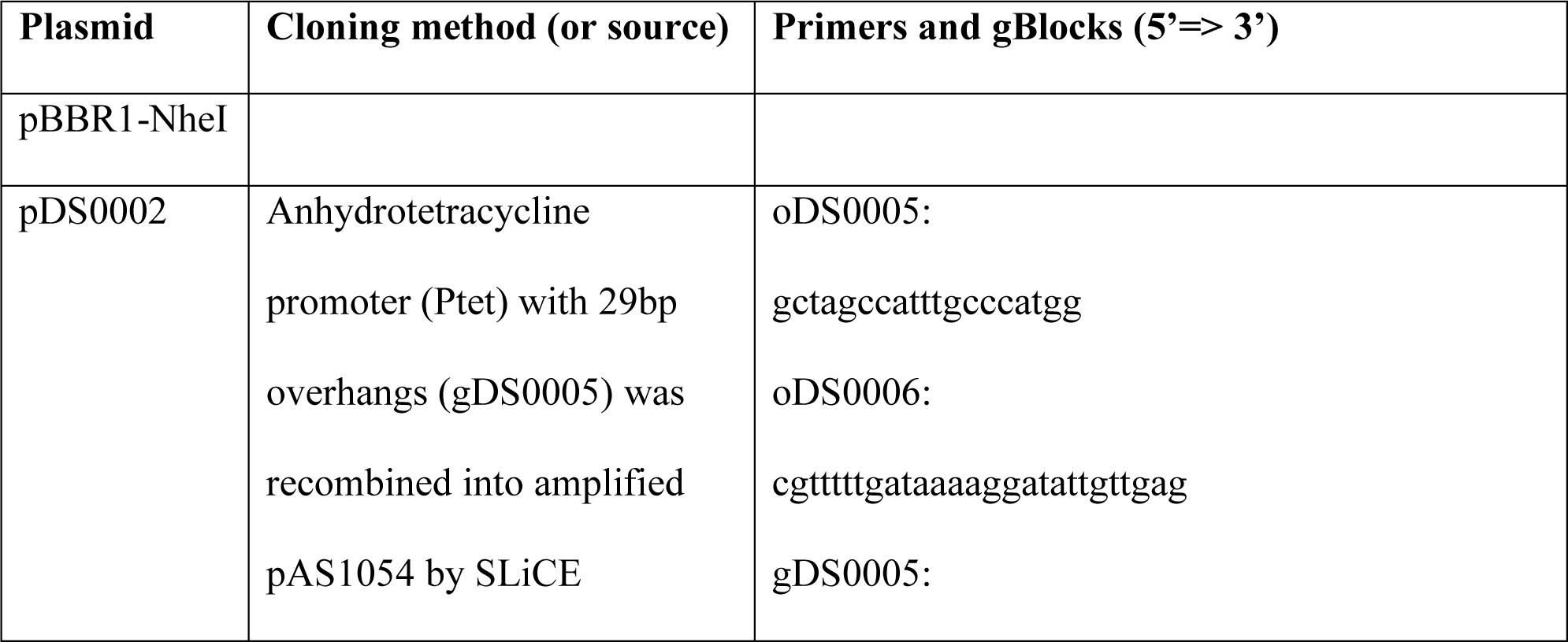
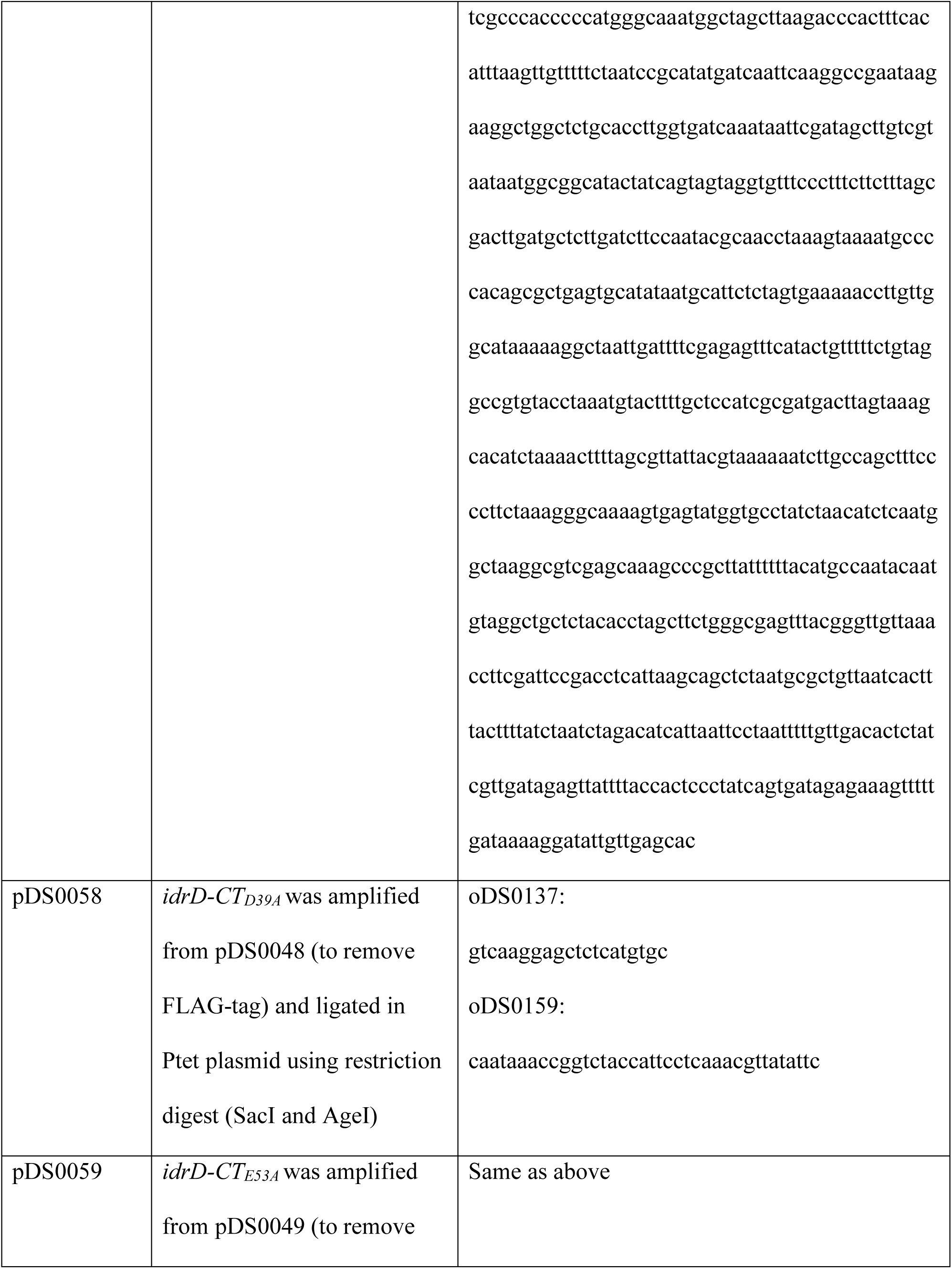
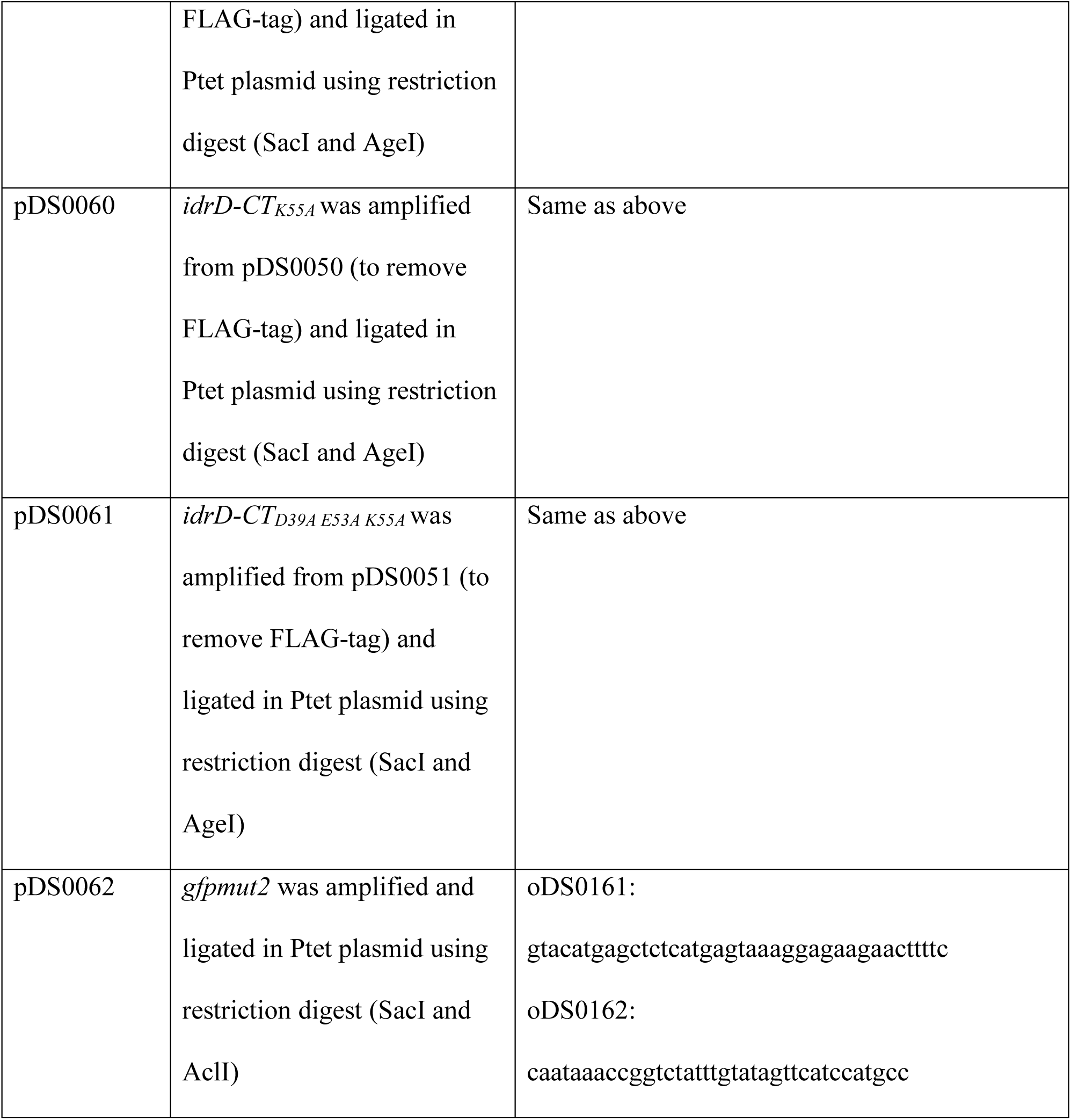
**Oligos used in this study.**

